# Pulse-driven and persistent antimicrobial resistance markers in a transboundary Great Lakes connecting channel: pulse-week-stratified water-quality thresholds for One Health surveillance

**DOI:** 10.64898/2026.06.26.734869

**Authors:** Xuexing Yao, Dennis Otieno, Qiudi Geng, Katelyn M. Brown, Lei Zhang, R. Michael McKay, Opeyemi U. Lawal

## Abstract

Surface waters in urban watersheds receive episodic inputs of wastewater, runoff, and road-salt residues during spring, yet the contribution of these short hydrological windows to antibiotic resistance gene (ARG) loading remains poorly resolved. Weekly samples were collected from offshore and nearshore sites in the Detroit River, a Great Lakes transboundary connecting channel, from February to December 2025. Five clinically relevant ARGs encoding resistance to carbapenems, methicillin, and colistin, alongside the fecal marker pepper mild mottle virus (PMMoV), were quantified by qPCR and paired with ten conventional water-quality variables. *blaNDM*, *mcr-1*, and *blaVIM-7* were not detected while *blaKPC* occurred as discrete pulses. One week (5 May) accounted for 37.2% of annual offshore *blaKPC* loading, and three weeks in late April to early May accounted for 72.8%, with peak concentrations reaching 5.4 × 10³ and 7.2 × 10³ copies L□¹. *mecA* was detected in nearly all samples without a dominant pulse. PMMoV normalization showed *blaKPC* did not vary seasonally (Kruskal-Wallis p = 0.198), consistent with diluted wastewater during spring precipitation events rather than an emergent source. *mecA*/PMMoV varied seasonally (p = 0.003) and was lowest in spring, implicating non-wastewater inputs in summer and autumn. Seven water-quality variables were significantly elevated during *blaKPC* pulse weeks. PCA distinguished pulse from background conditions, explaining 81.3% of variance. A random forest classifier achieved leave-one-out AUC of 0.917; ROC AUC reached 0.943 for total phosphorus, 0.924 for chloride, and 0.974 for the multivariate model. These results demonstrate that *blaKPC* and *mecA* operate through distinct source pathways and that routine water-quality monitoring can flag elevated *blaKPC* risk without additional sampling infrastructure.

## 1. Introduction

Antimicrobial resistance (AMR) is among the most serious public health challenges of the present century. Clinical surveillance has historically focused on hospitals, pharmacies, and food-production systems, but bacteria carrying resistance determinants also circulate through soil, sediment, and water (Berendonk et al., 2015; Larsson and Flach, 2022). Surface waters in urbanized catchments receive continuous inputs from wastewater treatment plants, combined sewer overflows, urban runoff, and agricultural drainage, and may act as both reservoirs and conduits for clinically important antibiotic resistance genes (ARGs) (Almakki et al., 2019; Pruden et al., 2021). Environmental ARG surveillance is therefore increasingly recognized as a necessary complement to clinical surveillance within a One Health framework (Hendriksen et al., 2019; McEwen and Collignon, 2018; Pruden et al., 2021).

Routine ARG monitoring in surface water, however, remains technically demanding. qPCR, high-throughput qPCR, metagenomic sequencing, and whole-genome sequencing can provide information on environmental AMR with high granularity, but their routine implementation requires specialized laboratory capacity, standardized workflows, quality-control procedures, and sustained funding (Cutrupi et al., 2024; Davis et al., 2023; Franklin et al., 2021; Liguori et al., 2022). These constraints have limited the adoption of ARG monitoring by water utilities and regulatory agencies outside of research settings. Much of the published surface-water AMR literature remains based on short-term or seasonally restricted sampling designs, which provide useful snapshots of ARG abundance but limited resolution of time-resolved dynamics (Alfahl et al., 2024; Bengtsson-Palme et al., 2023; Hart et al., 2023; Knapp et al., 2012). Relationships between ARGs and conventional water-quality variables may vary across seasons and hydrological conditions and can be obscured when these conditions are analyzed together. This limits the operational value of ARG measurements for routine risk assessment and highlights the need for analytical frameworks that relate ARG behavior to indicators already measured by government agencies (Huijbers et al., 2019; Liguori et al., 2022; Pruden et al., 2021).

North temperate urban watersheds are particularly challenging because spring rainfall and snowmelt can rapidly transport road-salt residues, accumulated particulates, nutrients, and microbial indicators over a few weeks, producing brief but intense pollutant pulses (Corsi et al., 2015; Galfi et al., 2016; Hrycik et al., 2024; Müller et al., 2020). Recent studies have increasingly examined associations between hydrological events and ARG dynamics. Rainfall and snowmelt were associated with shifts in bacterial communities and ARG concentrations in an urban river in Sapporo, Japan (Shayan et al., 2025), while substantial storm-driven ARG loads have been identified in urban streams (Garner et al., 2017). These studies suggested that short hydrological windows may contribute disproportionately to annual ARG variability. However, few studies have resolved these event-driven dynamics adopting sampling with high temporal resolution across an entire winter-to-spring transition.

The Detroit River is a Great Lakes transboundary connecting channel, part of the Erie-Huron corridor linking Lake Huron with Lake St. Clair and the western basin of Lake Erie, flowing through urbanized and historically industrial corridor (Burniston et al., 2018; Hartig et al., 2009; Scavia et al., 2019). The Detroit River corridor supports a binational urban population of approximately 5.5 million across the Windsor-Detroit metropolitan region (Hartig et al., 2009), and Lake Erie supplies drinking water to more than 12 million residents across Canada and the United States (Reutter, 2019). The system is managed under a long-standing binational water-quality framework, the Great Lakes Water Quality Agreement (Krantzberg, 2012; Scavia et al., 2019). Both sides of the Detroit River corridor include combined sewer infrastructure, and wet-weather overflows remain an important water-quality concern, especially in the nearshore (Hartig et al., 2021; Li et al., 2003). The River is a source of drinking water and supports recreational and provisioning fisheries along the Windsor and Detroit shorelines (Nguyen et al., 2025), all of which heighten its public health relevance. Combined with strong seasonal hydrological forcing from precipitation and runoff as well as existing monitoring infrastructure, these features make the Detroit River corridor a well-suited setting for AMR surveillance.

In this study, we report near-year-round weekly sampling of the Detroit River from February to December 2025 at one offshore and one nearshore site 5 clinically relevant ARGs were selected from the WHO Bacterial Priority Pathogens List (Sati et al., 2025) and the Canadian Antimicrobial Resistance Surveillance System (CARSS) (Public Health Agency of Canada, 2024). The three carbapenemase genes (*blaKPC*, *blaNDM*, *blaVIM-7*) hydrolyze carbapenems, a class reserved for serious multidrug-resistant Gram-negative infections. The carbapenem-resistant Enterobacterales that carry them are associated with high mortality and limited treatment options (Munoz-Price et al., 2013), *mecA* encodes the penicillin-binding protein PBP2a and confers resistance to nearly all β-lactams; although most often linked to methicillin-resistant *Staphylococcus aureus*, it is also common in coagulase-negative *staphylococci* (Chambers, 1997; Hiramatsu et al., 2013; Lakhundi and Zhang, 2018). *mcr-1* confers resistance to colistin, a last-line polymyxin, and was the first such resistance found to be carried on mobile plasmids (Liu et al., 2016).

This study had three objectives: (i) to characterize the occurrence and temporal behavior of these clinically relevant ARGs across an extended seasonal cycle, including the spring runoff window; (ii) to test whether year-round and pulse-week-stratified analyses provide different interpretations of ARG–water-quality relationships; and (iii) to evaluate whether routinely measured water-quality parameters can serve as practical early-warning indicators for periods of elevated AMR-relevant gene signal in a north temperate urban river. Since wastewater is a major source of ARGs in urban surface water, ARG concentrations were normalized to pepper mild mottle virus (PMMoV), a marker of human fecal pollution, to account for variation in wastewater input (Karkman et al., 2019; Kitajima et al., 2018; Rosario et al., 2009). ARG measurements were then compared with ten conventional water-quality variables collected at the offshore site through existing federal monitoring.

## 2. Materials and methods

### 2.1 Study area and sampling

Water samples were collected from two Canadian sites in the Detroit River near the Great Lakes Institute for Environmental Research in Windsor, Ontario, approximately 10 km downstream of Lake St. Clair. Both sites are part of the Detroit River node of the Environment and Climate Change Canada (ECCC) Connecting Channels Monitoring and Surveillance network. The offshore site (42.30815° N, 83.07475° W; ECCC station ON02GH0316) draws water from an intake >100 m from shore positioned near the riverbed at 11.2 m water depth, whereas the nearshore site (42.307621° N, 83.073746° W; ECCC station ON02GH0317), located approximately 10 m from shore, represents a shallower and more shoreline-influenced portion of the river (Fig. 1).

**Fig. 1.**
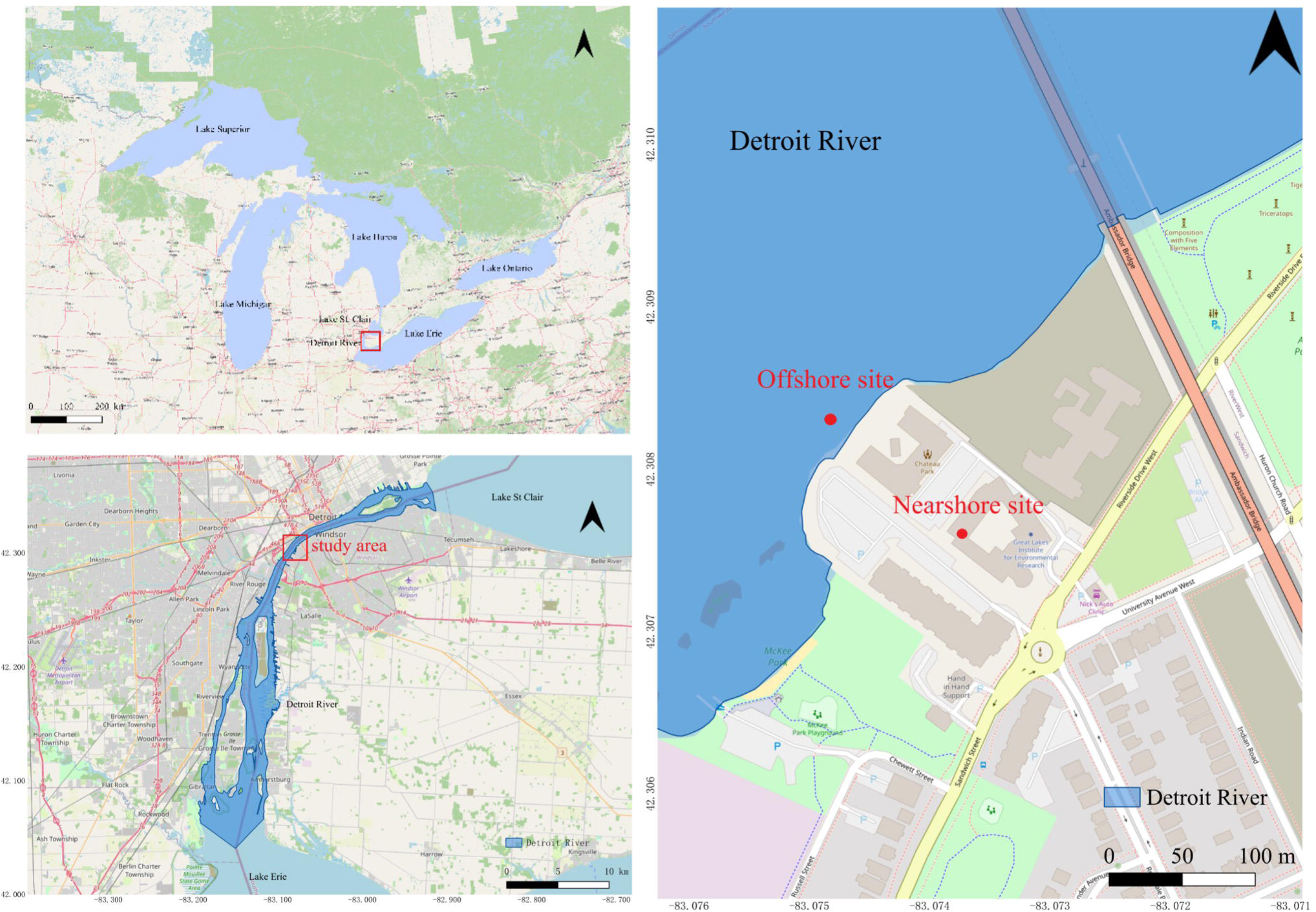
Study area and sampling locations in the Detroit River, a transboundary connecting channel between Ontario (Canada) and Michigan (USA). (a) Location of the Detroit River within the Laurentian Great Lakes system, linking Lake St. Clair to the western basin of Lake Erie. (b) The Detroit River showing the Canada–United States international border and the sampling area near Windsor, Ontario. (c) Enlarged view of the two sampling sites: the offshore site (ECCC station ON02GH0316, intake >100 m from shore at 11.2 m depth) and the nearshore site (ECCC station ON02GH0317, approximately 10 m from shore), both part of the Detroit River node of the ECCC Connecting Channels Monitoring and Surveillance network. Maps were produced in QGIS 3.44.

At each site, water was collected using a Teledyne ISCO 5800 refrigerated autosampler. The sampler collected approximately 155 mL every 4 h to generate 24 h composite samples in ProPak bottles, with 4 bottles collected per weekly sampling period for a total volume of approximately 4 L. For this study, samples collected between February and December 2025 were retained, yielding one weekly composite sample per site and covering all four seasons, including the spring runoff period. Samples were processed within 24 h of collection.

### 2.2 Water quality data

Water quality data were available only for the offshore site. The nearshore site samples included in this study did not have paired water quality measurements. Offshore site water quality data were obtained from the ECCC public dataset, accessed through the Open Government Portal (https://open.canada.ca/data/en/dataset/90e5f624-520a-4bd9-bedb-79b03c516a4d). Nine measured water-quality parameters were retrieved: turbidity (NTU), total phosphorus (TP), dissolved total phosphorus (DTP), soluble reactive phosphorus (SRP), dissolved organic carbon (DOC), filtered ammonia, nitrate plus nitrite (NO□□+ NO□□), total Kjeldahl nitrogen (TKN), and dissolved chloride. A derived particulate phosphorus ratio was calculated as PP ratio = (TP − DTP)/TP. Chlorophyll-*a* (Chl-*a*) was measured separately, as described below.

Samples were filtered, preserved, and shipped to ECCC’s National Laboratory for Environmental Testing (NLET) at the Canada Centre for Inland Waters (Burlington, ON), where they were analyzed according to standardized operating procedures. For the AMR-water analyses, offshore site AMR data were paired with water quality data by sampling date. AMR data from the nearshore site was used only for spatial comparisons of gene concentrations and were not included in water-quality-based statistical analyses.

Chl-*a* was measured by fluorometry. For each weekly composite sample, water was filtered through a Whatman GF/F filter at GLIER, and the filters were shipped to Bowling Green State University (BGSU). Pigments were extracted in 90% acetone buffered with magnesium carbonate, held in the dark for 24 h, and read on a Turner Designs TD-700 fluorometer using the non-acidification method (Welschmeyer, 1994). A 90% buffered acetone blank and a Turner Chl-*a* solid standard were used to calibrate the instrument prior to sample measurement. Chl-*a* concentrations (μg L^-1^) were calculated from the fluorometer reading, the extract volume, and the volume of water filtered.

### 2.3 Hydrological and meteorological data

Daily river discharge and meteorological data spanning the AMR sampling period were retrieved from public agency datasets (U.S. Geological Survey, 2025). Detroit River discharge was obtained from the United States Geological Survey (USGS) gauging station 04165710 (Detroit River at Fort Wayne at Detroit, MI; 42.301° N, 83.094° W). Daily mean discharge values originally reported in cubic feet per second were converted to cubic meters per second (1 cfs = 0.02832 m³ s□¹).

Daily precipitation and air temperature data were obtained from the ECCC Windsor Airport climate station (Climate ID 6139530; Station ID 54738; ICAO YQG; 42°16′34″ N, 82°57′19″ W), located approximately 10 km southeast of the offshore sampling site. Daily mean, maximum, and minimum air temperature and daily total precipitation were retrieved. A 7-day cumulative precipitation index was calculated as a rolling sum to summarize antecedent wet-weather conditions. Data were accessed from the ECCC historical climate data portal (https://climate.weather.gc.ca). Hydrological and meteorological variables were paired with the corresponding sampling date of each weekly composite sample.

### 2.4 Filtration and nucleic acid extraction

For each weekly sample, four 24 h composite bottles from a given site were combined to obtain a working volume of approximately 4 L. The combined sample was filtered through a Sterivex GP 0.22 μm capsule filter (Millipore, Burlington, MA, USA) using a peristaltic pump operated at approximately 200 ml min^-1^. Filtration continued until the full working volume had passed through the capsule or until the membrane clogged; therefore, the volume processed varied with sample turbidity. The filtered volume was recorded for each sample and used to express gene copy concentrations as copies L^-1^.

Total nucleic acid, including DNA and RNA, was co-extracted from each capsule using the AllPrep PowerViral DNA/RNA Kit (Qiagen, Germantown, MD, USA), following a previously described procedure (Harrop et al., 2026). In brief, the filter membrane was cut from the cartridge and bead-beaten in PM1 buffer with β-mercaptoethanol to lyse retained cells and viruses, and nucleic acid was recovered through the kit spin columns and eluted in 100 μL of RNase-free water. The same eluate served as template for DNA-based qPCR assays and the RNA-based RT-qPCR assay.

DNA concentration in the co-extracted nucleic acid eluate was quantified with a Qubit 4 fluorometer (Thermo Fisher Scientific, Waltham, MA, USA) using the dsDNA HS assay. Aliquots were stored at −80 °C and thawed immediately before analysis. A new Sterivex cartridge filter processed with sterile water served as an extraction blank in every batch and remained below detection in all qPCR and RT-qPCR assays.

### 2.5 qPCR and RT-qPCR quantification

Probe-based qPCR and RT-qPCR assays were used to quantify six molecular markers, including five ARGs, *blaKPC*, *blaNDM*, *blaVIM-7*, *mecA*, and *mcr-1*, as well as the wastewater-associated fecal marker pepper mild mottle virus (PMMoV). PMMoV was included as a wastewater-associated human fecal marker because it is abundant in human wastewater, widely used in microbial water-quality studies, and relatively persistent in aquatic environments (Kitajima et al., 2018; Rosario et al., 2009; Symonds et al., 2019). Normalizing ARG concentrations to PMMoV separates changes in wastewater loading from changes in the resistance signal, because fecal contamination is a primary driver of ARG abundance in impacted surface waters (Karkman et al., 2019). Deviations from this marker then flag ARG inputs that do not track wastewater input. PMMoV has been used in this way to normalize carbapenemase-gene measurements in the same wastewater system (Harrop et al., 2026). The triplex assay quantified *blaKPC*, *blaNDM*, and *blaVIM-7*, using primer-probe sets reported previously for these carbapenemase genes (Brown-Jaque et al., 2018; Hindiyeh et al., 2008; Lee et al., 2015), Primer-probe sets for these assays were based on published primer-probe sets, as listed in Table S1.

DNA-based qPCR assays were performed using Luna Universal Probe qPCR Master Mix on an MA-6000 real-time PCR system (Aumintec, Richmond Hill, ON, Canada). The *blaKPC*, *blaNDM*, and *blaVIM-7* assay was run as a triplex reaction, whereas *mecA* and *mcr-1* were quantified in singleplex reactions. PMMoV RNA was measured by one-step RT-qPCR using the Luna Probe-based one-step RT-qPCR system (New England Biolabs, Ipswich, MA, USA). Reaction setup, thermal cycling conditions, and six-point gBlock standard curves are provided in Table S2.

All samples were analyzed in technical triplicate with no-template controls, extraction blanks, and six-point gBlock standard curves on each plate. Standard curves met MIQE-recommended performance criteria, with efficiencies of 90–110% and R² ≥ 0.99 (Bustin et al., 2009). The limit of detection (LOD) and limit of quantification (LOQ) for each assay were determined from replicate standard curves, with the LOD was defined as the lowest concentration detected with 95% probability, and the LOQ as the lowest concentration quantifiable at a 35% coefficient of variation (Harrop et al., 2026). Samples below the assay LOD were reported as non-detects.

### 2.6 Statistical analysis

Statistical analyses were conducted in Python 3.11 using NumPy, Pandas, SciPy, and scikit-learn (Pedregosa, 2011). Spatial maps were prepared in QGIS 3.44, and selected figures were produced in Origin 2021. Seasons were defined as winter (December–February), spring (March–May), summer (June–August), and autumn (September–November). Results were considered statistically significant at p < 0.05, or adjusted p < 0.05 for multiple comparisons.

#### 2.6.1 Temporal and pulse-week-stratified analyses

Weekly AMR gene concentrations were ranked to estimate the amount of the annual signal that was contributed by short duration, high-concentration periods. The cumulative contribution of the highest 1, 3, and 5 weeks was calculated for each detected ARG marker. To account for wastewater input, *blaKPC* and *mecA* were normalized to PMMoV, and seasonal differences in these ratios were tested using Kruskal–Wallis tests (Benjamini and Hochberg, 1995).

Spring *blaKPC* pulse weeks were defined as spring weeks when offshore *blaKPC* exceeded the annual median concentration of 186 copies L^-1^. This identified four pulse weeks. Summer and autumn weeks were used as the background group (n = 21). Winter was excluded because several of the samples from this period exceeded the 186 copies L^-1^ threshold and chloride concentrations were elevated, likely reflecting road de-icing inputs. Differences in water-quality variables between pulse weeks and background weeks were tested using Mann-Whitney U test, with effect sizes reported as absolute rank-biserial correlations, |r_B|.

#### 2.6.2 Multivariate analysis

Principal component analysis (PCA) was used to summarize variation in the 10 water-quality variables across sampling weeks with complete observations (n = 41). Variables were centered and scaled to unit variance before PCA (Jolliffe and Cadima, 2016). The first two principal components were plotted to visualize seasonal clustering, with 95% confidence ellipses drawn for each season.

To test whether spring weeks could be distinguished from the rest of the year, we trained a random forest classifier using the same 10 water-quality variables as predictors (Breiman, 2001). The model was implemented in scikit-learn with 500 trees and evaluated by leave-one-out cross-validation (Pedregosa, 2011). Predictor importance was estimated using Gini-based importance and verified with permutation importance. Partial dependence plots were produced for the four most important variables to examine possible threshold-like responses.

#### 2.6.3 ROC analysis and threshold identification

Receiver operating characteristic (ROC) analysis was used to assess how well the seven water-quality variables identified in the univariate comparisons distinguished spring weeks (n = 8) from non-spring weeks (n = 24), using the same spring vs non-spring grouping as the random forest classifier in Section 2.6.2. Both groups were restricted to weeks with a paired *blaKPC* measurement and complete water-quality data, which retained 8 of the 12 spring weeks and 24 non-spring weeks. The classification analyses used spring versus non-spring grouping rather than the *blaKPC* > 186 copies L[¹ threshold used in the pulse-week comparison. Spring concentrated the observed *blaKPC* pulses, and the classifier was intended to flag periods of elevated risk rather than to compare water quality within already-identified pulse weeks. AUC values and 95% confidence intervals were estimated from 2,000 stratified bootstrap resamples (Efron, 1979; Hanley and McNeil, 1982). Optimal thresholds were defined by maximizing Youden’s statistic (*J* = sensitivity + specificity – 1) on the empirical ROC curve (Youden, 1950). A logistic regression model including all seven variables was also evaluated using the same procedure.

## 3. Results

### 3.1 Temporal dynamics of *blaKPC*, *mecA*, and PMMoV in the Detroit River

Six genetic markers, including: *blaKPC*, *mecA*, *blaNDM*, *mcr-1*, *blaVIM-7*, and PMMoV, were analyzed in this study. *blaNDM*, *mcr-1*, and *blaVIM-7* remained below detection limits across all sampling weeks and both sites throughout the year and are therefore not discussed further. The three consistently detected markers: *blaKPC* and *mecA* as clinically relevant resistance genes, and PMMoV as a human fecal wastewater tracer were assessed.

Weekly measurements from February to December 2025 showed clearly different temporal patterns for the three markers at the offshore and nearshore site (Fig. 2). Offshore *blaKPC* remained near or below detection limits for most of the year but rose sharply during spring to 5,414 copies L^-1^ on 28 April and 7,242 copies L^-1^ on 5 May 2025. Outside these two weeks, offshore *blaKPC* rarely exceeded 700 copies L^-1^.

**Fig. 2.**
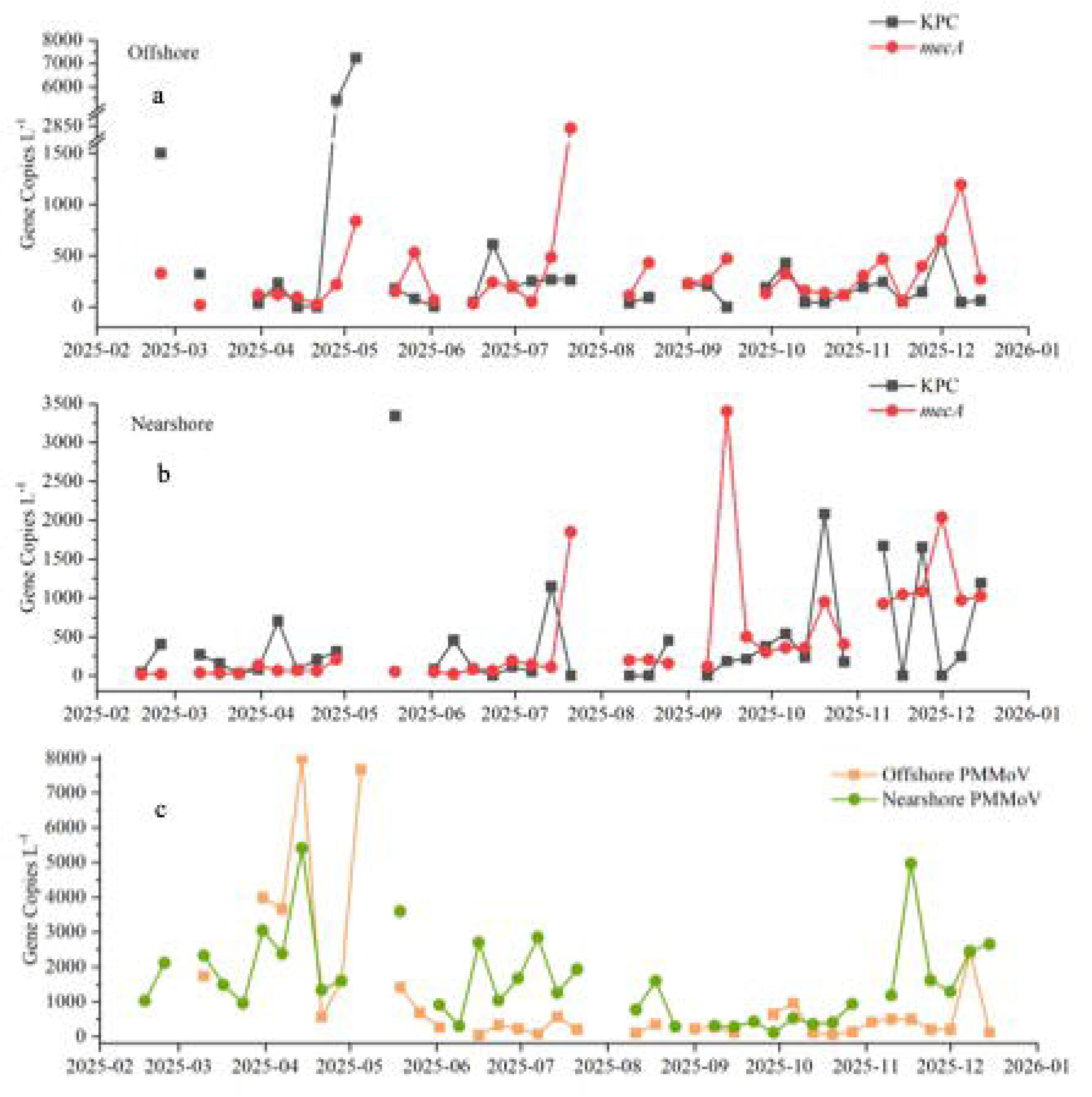
Temporal dynamics of *blaKPC*, *mecA*, and PMMoV in the Detroit River, February to December 2025. Weekly gene copy concentrations (copies L^-1^) are shown for (a) *blaKPC* and *mecA* at the offshore site, (b) *blaKPC* and *mecA* at the nearshore site, and (c) PMMoV at both sites. *blaKPC* occurred as short spring pulses, whereas *mecA* was detected in nearly all weeks.

In contrast*, mecA* was detected in nearly every weekly sample at both sites. Offshore concentrations mostly ranged from tens to several hundred copies L^-1^, with the highest value of 2,825 copies L^-1^ recorded on 21 July and a secondary peak of 1,194 copies L^-1^ in early December. Unlike *blaKPC*, *mecA* showed no single dominant peak and was detected throughout the year.

PMMoV was detected throughout the year at both sites and rose to its highest single-week values in late winter and spring (Fig. 2c). Offshore PMMoV reached 32,006 copies L^-1^ on 24 February and of 7,681 copies L^-1^ on 5 May, the same week as the highest *blaKPC* value. The May PMMoV secondary peak coincided closely with offshore *blaKPC* maximum.

Spatial comparison showed that offshore *blaKPC* during spring was substantially higher than nearshore *blaKPC*, whose maximum was 3,346 copies L^-1^ on 19 May. The pattern reversed for *mecA* in autumn: nearshore *mecA* reached 3,407 copies L^-1^ on 15 September and stayed above 900 copies L^-1^ for several subsequent weeks, while offshore *mecA* during the same period remained below 500 copies L^-1^.

### 3.2 Quantitative characterization of AMR gene dynamics

To quantify the temporal distribution of AMR gene loads, we calculated each peak week’s contribution to the annual sum of weekly concentrations (Fig. 3a, b). Of the annual sum of weekly *blaKPC* concentrations at offshore site, the single highest week (7,242 copies L□¹ on 5 May) accounted for 37.2%. The top three weeks together contributed 72.8%, and the top five weeks contributed 79.3%. The top week of *mecA* at the offshore site contributed 23.4% of its annual sum, and the top three weeks contributed 40.3%.

**Fig. 3.**
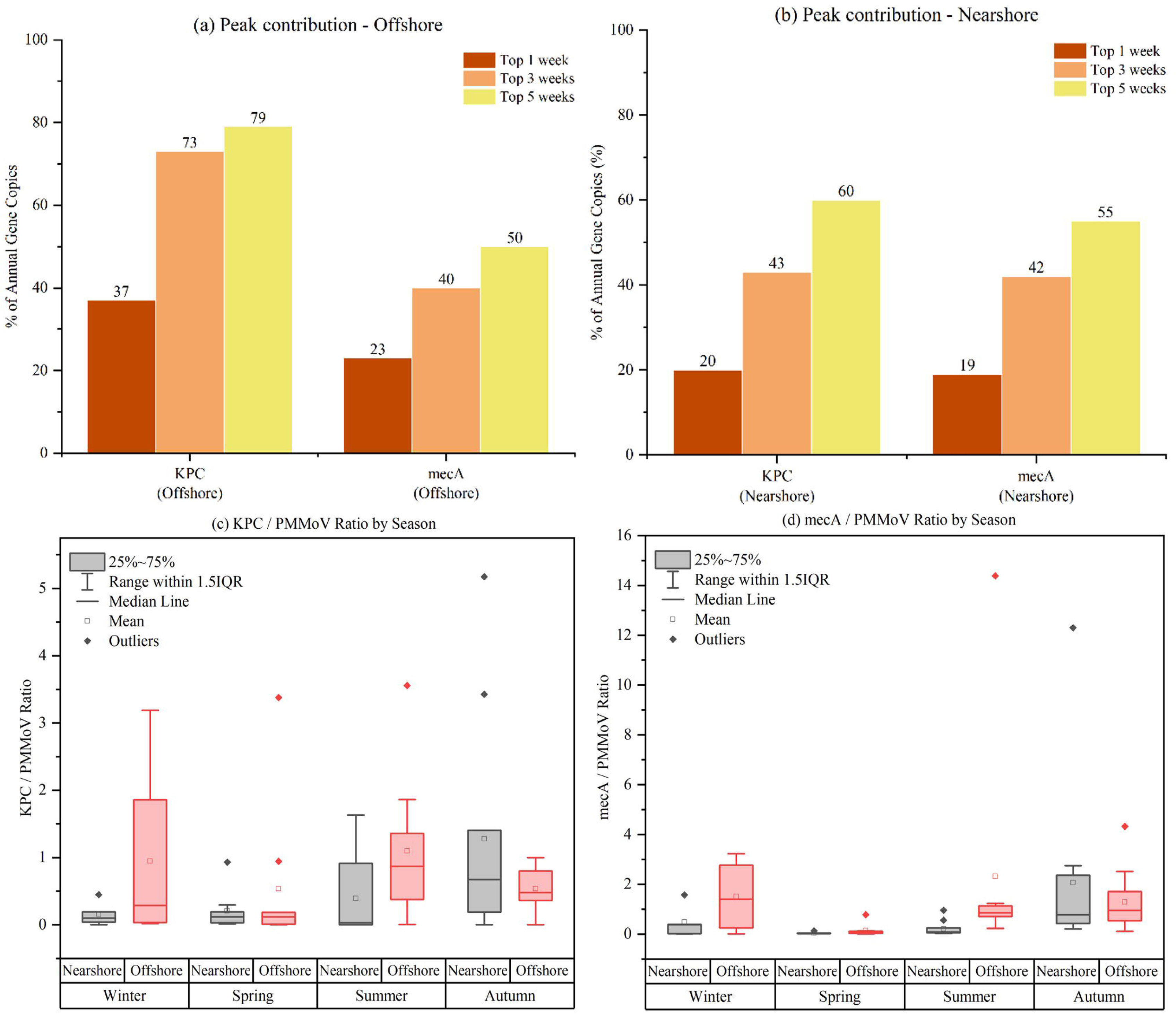
Quantitative characterization of *blaKPC* and *mecA* dynamics in the Detroit River, 2025. (a, b) Cumulative contribution of the highest-concentration weeks to the annual sum of weekly gene concentrations for *blaKPC* and *mecA* at the offshore and nearshore sites; (c, d) seasonal variation in the *blaKPC*/PMMoV and *mecA*/PMMoV ratios at both sites.

At the nearshore site, the top week of *blaKPC* contributed 20.2% of its annual sum, and 19.8% of the *mecA* annual sum. The top three weeks contributed 42.8% for nearshore *blaKPC* and 42.3% for nearshore *mecA*.

To examine the relationship between AMR genes and wastewater input across seasons, we normalized *blaKPC* and *mecA* concentrations to PMMoV at each sampling time (Fig. 3c, d). The *blaKPC*/PMMoV ratio at the offshore site did not differ significantly among seasons (Kruskal–Wallis p = 0.198), with median values of 0.29 in winter, 0.12 in spring, 0.87 in summer, and 0.48 in autumn.

The *mecA*/PMMoV ratio, by contrast, showed clear seasonal variation (Fig. 3d). Offshore median ratios were 1.40 in winter, 0.04 in spring, 0.86 in summer, and 0.95 in autumn (Kruskal–Wallis p = 0.003). Pairwise comparisons showed that the spring median was significantly lower than those of summer (Mann–Whitney p = 0.002) and autumn (p = 0.001). When PMMoV was highest (spring), *mecA*/PMMoV was lowest; when PMMoV was lower (summer and autumn), *mecA*/PMMoV was highest.

### 3.3 Hydrological forcing and seasonal water-quality patterns

Daily Detroit River discharge remained close to 5,687 ± 243 m³ s□¹ from March through November 2025, with no pronounced spring peak (Fig. S1b). By contrast, daily precipitation showed several wet-weather events throughout the same period (Fig. S1a). The largest single-day rainfall of the year was 40.3 mm on 2 April, after which 7-day cumulative precipitation reached the annual peak of 56.2 mm. A second multi-day event during 4-6 May produced a 7-day cumulative total 36.7 mm on 5 May. Additional daily totals above 20 mm occurred on 4 June, 18 June, and 9 July. Outside these episodes, daily precipitation generally remained below 10 mm.

Weekly water quality monitoring at the offshore site captured clear seasonal variation across the five major groups of parameters examined: turbidity and chloride, phosphorus fractions, dissolved inorganic nitrogen, and total Kjeldahl nitrogen with dissolved organic carbon (Fig. 4). Turbidity remained below 10 NTU for most of the year, but rose markedly during spring, reaching 75.3 NTU on 21 April and 101 NTU on 5 May 2025 (Fig. 4a). Chloride showed a similar increase in spring, with the annual maximum of 33.2 mg L^-1^ recorded on 7 April. Both parameters returned to low levels by early June and remained stable through summer and autumn, with summer and autumn medians of approximately 5.6 NTU and 9.4 mg L^-1^, respectively. Phosphorus fractions followed the same temporal pattern (Fig. 4b). TP peaked at 0.111 mg L^-1^ on 5 May, compared with summer and autumn values near 0.010 mg L^-1^. DTP reached a maximum of 0.028 mg L^-1^ on 7 April, and SRP peaked at 0.022 mg L^-1^ on 14 April. Both DTP and SRP were substantially higher than their summer and autumn values of approximately 0.002 mg L^-1^ and 0.0004 mg L^-1^, respectively.

**Fig. 4.**
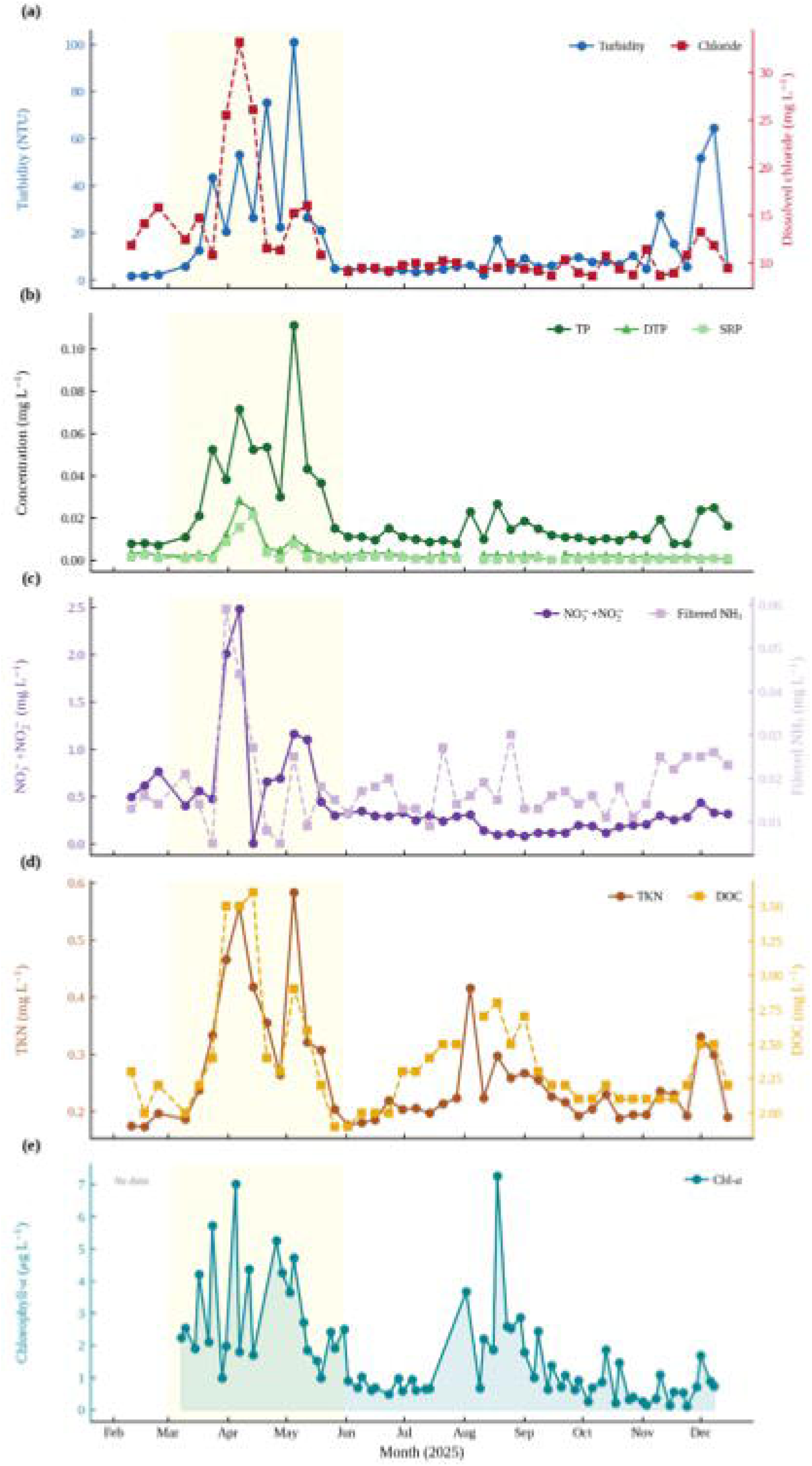
Seasonal variation in offshore water-quality parameters in the Detroit River, 2025. (a) Turbidity and dissolved chloride; (b) phosphorus fractions (total phosphorus, TP; dissolved total phosphorus, DTP; soluble reactive phosphorus, SRP); (c) dissolved inorganic nitrogen (NO□□+NO□□and filtered NH□); (d) total Kjeldahl nitrogen (TKN) and dissolved organic carbon (DOC); (e) Chl-*a*. Most parameters reached annual maxima in spring.

Dissolved inorganic nitrogen species showed different dynamics (Fig. 4c). NO□□+ NO□□reached its annual maximum of 2.48 mg L^-1^ on 7 April and remained elevated through early May, then declined to summer and autumn values near 0.22 mg L^-1^. Filtered NH□showed smaller seasonal variation, with values generally between 0.010 and 0.030 mg L^-1^ throughout the year and no pronounced spring peak. TKN and DOC were less variable across seasons (Fig. 4d). TKN ranged from 0.17 to 0.58 mg L^-1^, with slightly elevated values during spring (median 0.41 mg L^-1^) but considerable overlap with other seasons (summer and autumn median 0.21 mg L^-1^). DOC values ranged from 1.9 to 3.6 mg L^-1^ with only minor seasonal change. Chlorophyll-*a* at the offshore site rose during spring, reaching 7.01 μg L□¹ on 5 April with elevated values continuing through May (Fig. 4e). A secondary peak of 7.26 μg L□¹ occurred on 18 August. The spring median was 2.41 μg L□¹, with summer and autumn medians of 0.91 and 0.66 μg L□¹, respectively. Multiple water-quality parameters therefore reached their annual maxima within the same spring period in which offshore *blaKPC* peaked and major precipitation events occurred (Figs. 2a, 4, S1).

### 3.4 Water-quality conditions associated with spring *blaKPC* pulse weeks

To identify the water-quality conditions associated with the spring *blaKPC* pulses observed in this study (see Fig. 2), we compared water quality during spring weeks with elevated *blaKPC* to water quality during the summer and autumn background period. Spring *blaKPC* pulse weeks were defined as spring weeks with *blaKPC* above the annual median (186 copies L^-1^; n = 4). Summer and autumn weeks (n = 21) formed the background group; winter weeks were excluded for reasons described in Section 2.6.1. The two groups were compared using Mann-Whitney U tests for each of ten water-quality parameters (Fig. 5, Table 1).

**Fig. 5.**
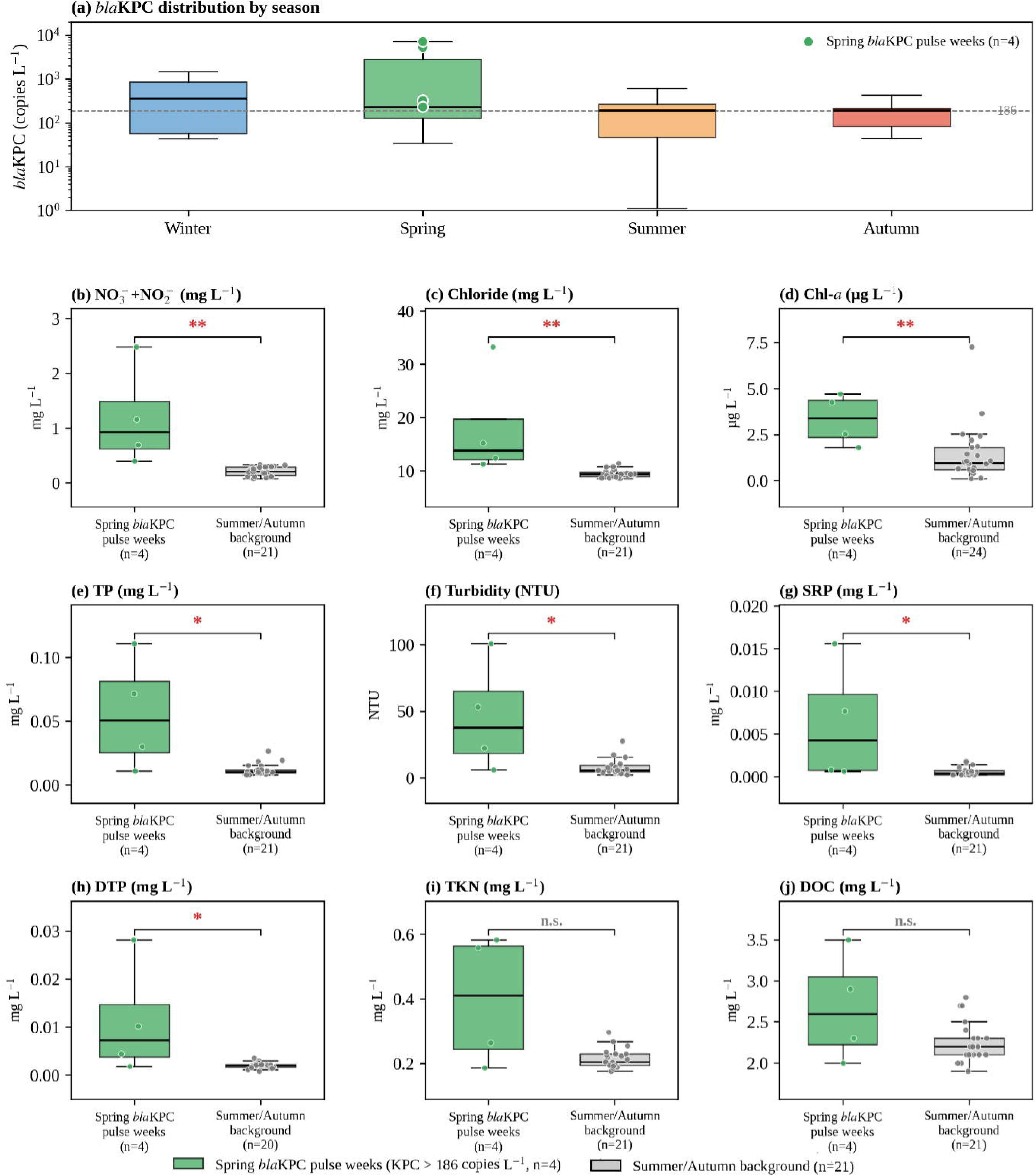
Water-quality conditions during spring *blaKPC* pulse weeks compared with the summer–autumn background in the Detroit River, 2025. (a) Seasonal distribution of offshore *blaKPC* on a logarithmic scale. Filled green circles mark the four spring pulse weeks used in the analysis, and the dashed line indicates the 186 copies L□¹ pulse threshold. Non-detects are not shown. (b–j) Mann–Whitney U comparisons of nine water-quality variables between spring *blaKPC* pulse weeks (n = 4) and the summer–autumn background, ordered by decreasing significance. Pulse weeks were spring weeks with offshore *blaKPC* above the annual median of 186 copies L□¹. The background comprises summer and autumn weeks with measured *blaKPC* (n = 21), except for Chl-*a* (all matched summer–autumn weeks, n = 24) and DTP (n = 20, one week without a measurement). Boxes show the median and interquartile range, and overlaid points are individual weekly values. Brackets give the pairwise test result: *p < 0.05, **p < 0.01; n.s., not significant.

**Table 1.** Water quality during spring *blaKPC* pulse weeks and background periods in the Detroit River, 2025 Note: Values are medians. Spring events were defined by *blaKPC* > 186 copies LO¹; background includes summer and autumn samples. p-values are from Mann–Whitney *U* tests. | r_B | = absolute rank-biserial effect size. * p < 0.05, ** p < 0.01, *** p < 0.001; n.s., not significant. Spring blaKPC pulse weeks n = 4, background n = 21, except Chl-*a* (n = 24) and DTP (n = 20).

| Variable | Unit | Spring<br>*blaKPC* pulse<br>weeks<br>Median | Background<br>Median | p-value | \|r<sub>B</sub>\| | Sig. |
| --- | --- | --- | --- | --- | --- | --- |
| NO<sub>3</sub><sup>-</sup>+NO<sub>2</sub><sup>-</sup> | mg<br>L<sup>-1</sup> | 0.93 | 0.21 | 0.002 | 1.00 | ** |
| Chloride | mg<br>L<sup>-1</sup> | 13.8 | 9.35 | 0.003 | 0.98 | ** |
| Chlorophyll-*a* | µg L<sup>-1</sup> | 3.40 | 0.964 | 0.009 | 0.79 | ** |
| Total phosphorus<br>(TP) | mg<br>L<sup>-1</sup> | 0.051 | 0.010 | 0.016 | 0.79 | * |
| Turbidity | NTU | 37.8 | 5.63 | 0.019 | 0.74 | * |
| Soluble reactive<br>phosphorus (SRP) | mg<br>L<sup>-1</sup> | 0.0043 | 0.0004 | 0.019 | 0.74 | * |
| Dissolved total<br>phosphorus (DTP) | mg<br>L<sup>-1</sup> | 0.0073 | 0.0020 | 0.044 | 0.66 | * |
| Total Kjeldahl<br>nitrogen (TKN) | mg<br>L<sup>-1</sup> | 0.41 | 0.21 | 0.128 | 0.50 | n.s. |
| Dissolved organic<br>carbon (DOC) | mg<br>L<sup>-1</sup> | 2.60 | 2.20 | 0.229 | 0.39 | n.s. |
| Filtered NH<sub>3</sub> | mg<br>L<sup>-1</sup> | 0.023 | 0.015 | 0.281 | 0.36 | n.s. |

**Note:** Values are medians. Spring events were defined by *blaKPC* > 186 copies L<sup>-1</sup>; background includes summer and autumn samples. p-values are from Mann–Whitney *U* tests. |r<sub>B</sub>| = absolute rank-biserial effect size. \* p < 0.05, \*\* p < 0.01, \*\*\* p < 0.001; n.s., not significant. Spring *blaKPC* pulse weeks n = 4, background n = 21, except Chl-*a* (n = 24) and DTP (n = 20).

Seven of the ten parameters were significantly higher during spring *blaKPC* pulse weeks than during the summer and autumn background (Fig. 5b-h). NO□□+ NO□□showed the largest difference, with a spring median of 0.93 mg L^-1^ compared with 0.21 mg L^-1^ during background weeks (p = 0.002, |r_B| = 1.00). Chloride was also strongly elevated during these pulse weeks, with medians of 13.8 mg L^-1^ and 9.35 mg L^-1^ respectively (p = 0.003, |r_B| = 0.98). The rank-biserial effect sizes of 1.00 and 0.98 mean that every spring *blaKPC* pulse week had higher NO□□+ NO□□and higher chloride than almost every summer or autumn background week.

TP and turbidity were also significantly higher during spring pulse weeks. TP medians were 0.051 mg L^-1^ during pulse weeks and 0.010 mg L^-1^ during the background (p = 0.016, |r_B| = 0.79). Chl-*a* medians were 3.40 μg L□¹ during pulse weeks and 0.964 μg L□¹ during the background (p = 0.009, |r_B| = 0.79). Turbidity medians were 37.8 NTU and 5.63 NTU (p = 0.019, |r_B| = 0.74). SRP and DTP showed smaller but still significant differences: SRP medians were 0.0043 mg L^-1^ during pulse weeks and 0.0004 mg L^-1^ during the background (p = 0.019, |r_B| = 0.74), and DTP medians were 0.0073 mg L^-1^ and 0.0020 mg L^-1^ (p = 0.044, |r_B| = 0.66). The remaining three parameters, TKN, DOC, and filtered NH□, did not differ significantly between groups (Fig. 5i-j, Table 1).

### 3.5 Multivariate analysis of water quality and AMR dynamics

#### 3.5.1 Principal component analysis

A PCA was performed on the ten standardized water-quality variables across all sampling weeks with complete data (n = 41) to examine whether overall water quality structure varied by season and how it related to *blaKPC* and *mecA* concentrations. PC1 explained 62.4% of the total variance and the PC2 explained 18.9%, giving a cumulative 81.3% for two-dimensional biplot (Fig. 6).

**Fig. 6.**
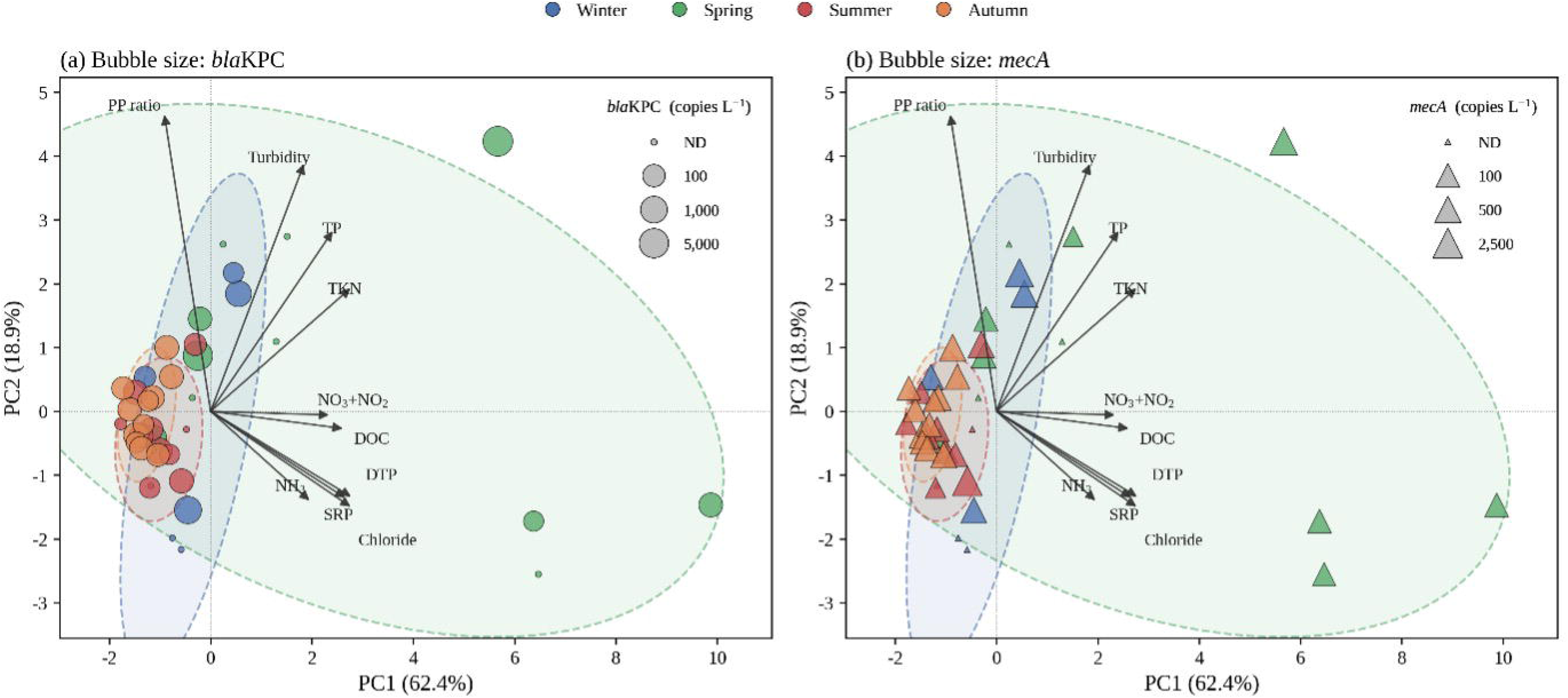
Principal component analysis (PCA) of ten offshore water-quality variables in the Detroit River (2025; n = 41 weeks). PC1 and PC2 explained 62.4% and 18.9% of the variance. Arrows show variable loadings, points are colored by season, and dashed ellipses give 95% confidence regions. Marker size is proportional to log□gene concentration, with non-detects (ND) at the smallest size: (a) circles sized by *blaKPC*, (b) triangles sized by *mecA*. Both panels share the same ordination and differ only in the gene by scaling the markers.

All ten water-quality variables loaded positively on PC1, with largest contributions from DTP (0.370), TKN (0.369), chloride (0.369), SRP (0.355), and DOC (0.348). PP ratio had a small negative loading (−0.122) but no other variable loaded strongly in the negative direction. PC2 was dominated by PP ratio (+0.620) and turbidity (+0.517) in the positive direction, and NH□(−0.187) and chloride (−0.200) in the negative direction.

Samples were separated clearly by season along PC1 (Fig. 6). Mean PC1 scores were +2.68 for spring, −1.26 for autumn, −1.02 for summer, and −0.35 for winter. Confidence ellipses constructed at two standard deviations showed that the spring cluster did not overlap with the summer or autumn clusters along PC1. The three spring samples with the highest *blaKPC* concentrations occupied the most positive PC1 (Fig. 6a), with marker sizes larger than those of any summer or autumn sample. When markers were sized by *mecA* instead, the largest values fell mainly within the summer and autumn cluster near the origin rather than in the positive-PC1 spring region (Fig. 6b).

#### 3.5.2 Random forest classification and partial dependence

A random forest classifier was trained to distinguish spring weeks from non-spring weeks using the ten water-quality variables as predictors. Leave-one-out cross-validation gave an area under the ROC curve of 0.917 (Fig. 7a).

**Fig. 7.**
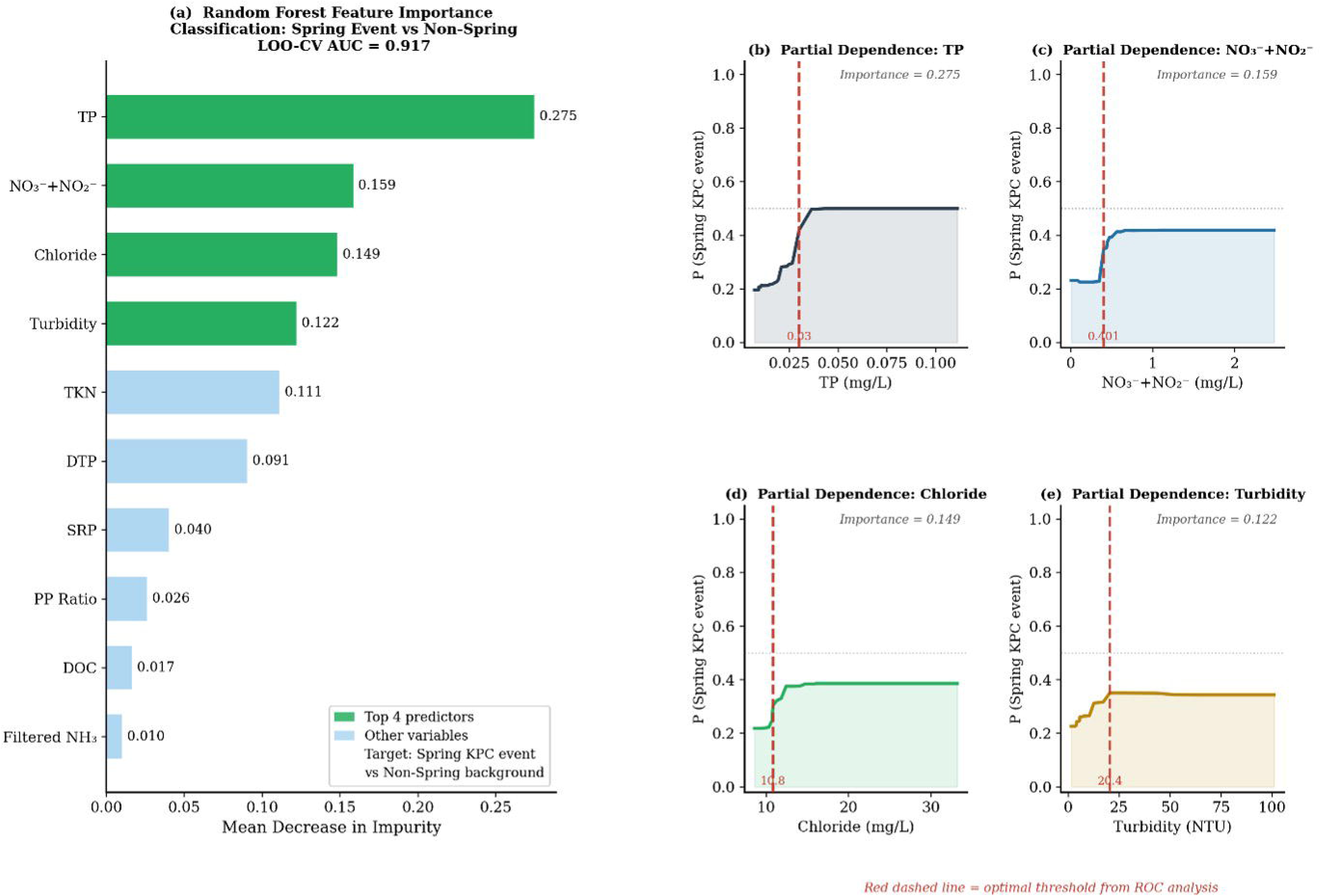
Random Forest classification of spring weeks (n = 8) versus non-spring weeks (n = 24) from ten offshore water-quality variables in the Detroit River, 2025. (a) Gini-based feature importance; the classifier reached a leave-one-out cross-validation (LOO-CV) AUC of 0.917. (b)–(e) Partial dependence plots for the four most important predictors, showing non-linear threshold responses. Red dashed lines mark the ROC-derived optimal warning thresholds.

Feature importance ranked TP highest (0.247), followed by NO□□+ NO□□(0.186), chloride (0.149), turbidity (0.122), and TKN (0.111). The remaining five variables each had importance values below 0.11 (Fig. 7a). The same four top variables were identified by permutation importance, with identical ranking.

Partial dependence plots for the four top variables showed threshold-type responses rather than linear relationships (Fig. 7b-e). The predicted probability of a spring week remained near 0.20 when TP was below approximately 0.030 mg L^-1^, then increased sharply to around 0.50 at higher TP values. Similar step changes occurred for NO□□+ NO□□around 0.40 mg L^-1^, chloride around 10.8 mg L^-1^, and turbidity around 20 NTU. The inflection points of the partial dependence curves were close to the optimal decision threshold obtained from the ROC analysis.

Year-round Spearman correlations between *blaKPC* and each individual water-quality variable were calculated for comparison. None of the ten correlations was significant. All had |ρ| below 0.30, and the lowest p-value among them was 0.089.

### 3.6 Water-quality thresholds for flagging spring *blaKPC* pulse periods

ROC analysis was performed for the seven water-quality variables identified in Section 3.4, to assess their ability to discriminate spring weeks (n = 8) from non-spring weeks (n = 24) as a single-variable classifier (Fig. 8, Table 2).

**Fig. 8.**
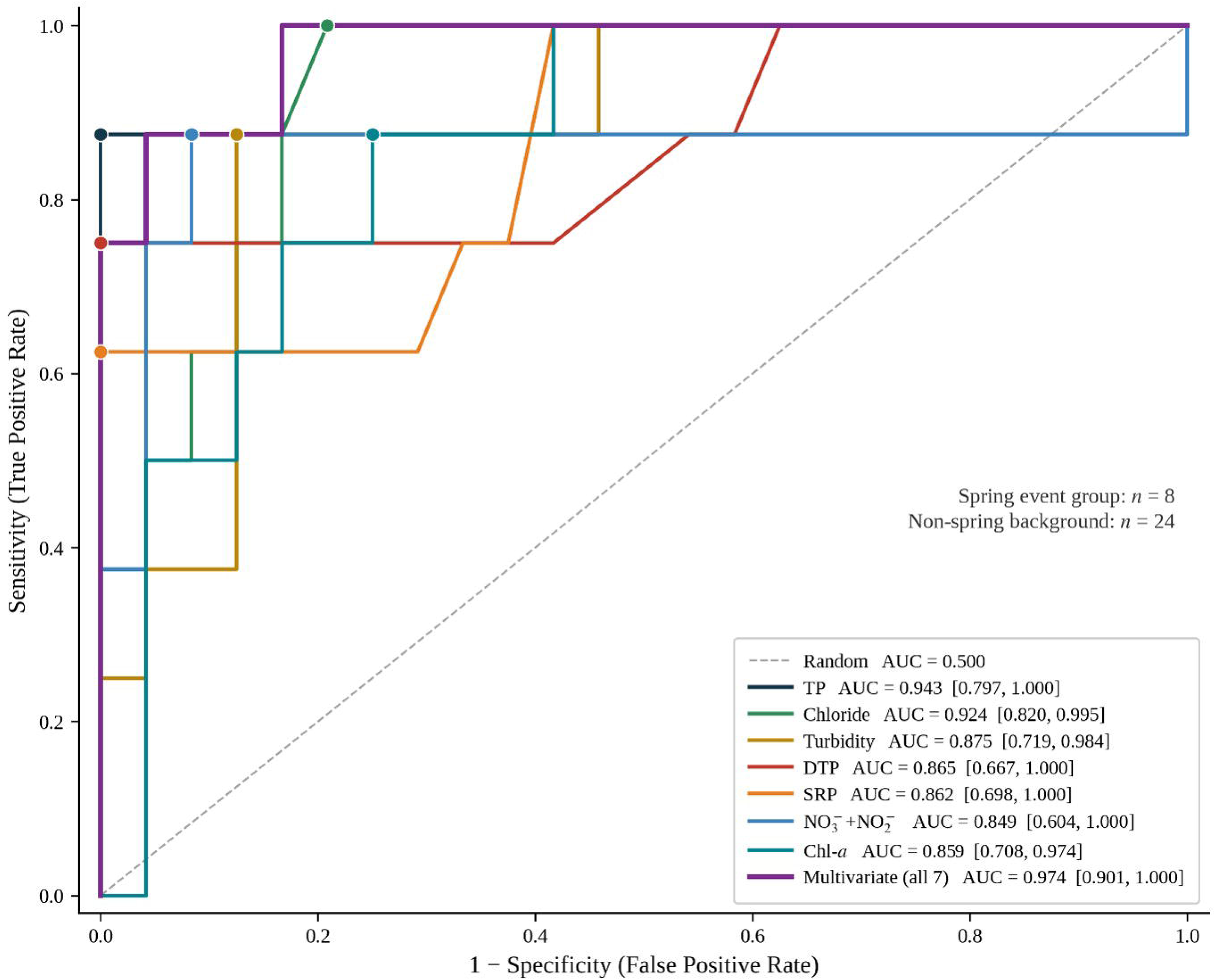
Receiver operating characteristic (ROC) analysis of seven water quality variables as single-variable predictors of spring weeks (n = 8) versus non-spring weeks (n = 24) in the Detroit River, 2025. Area under the curve (AUC) values with 95% bootstrap confidence intervals (2000 resamples) are shown in the legend. Filled circles mark Youden’s J optimal thresholds.

**Table 2.** Receiver operating characteristic (ROC) analysis of water quality variables as predictors of spring weeks in the Detroit River, 2025. Note: AUC = area under the ROC curve. Confidence intervals were estimated using 2,000 bootstrap resamples. Optimal thresholds were selected by Youden’s *J* statistics. Spring weeks, n = 8; non-spring weeks, n = 24. The classification grouping (spring vs non-spring) differs from the univariate pulse-week comparison in Table 1 (n = 4 spring *blaKPC* pulse weeks vs n = 21 summer–autumn background); see Methods 2.6.1 and 2.6.3.

| Variable | AUC | 95% CI | Optimal threshold | Sensitivity | Specificity |
| --- | --- | --- | --- | --- | --- |
| TP | 0.943 | [0.803, 1.000] | > 0.030 mg L<sup>-1</sup> | 0.88 | 1.00 |
| Chloride | 0.924 | [0.821, 0.995] | > 10.8 mg L<sup>-1</sup> | 1.00 | 0.79 |
| Turbidity | 0.875 | [0.724, 0.983] | > 20.4 NTU | 0.88 | 0.88 |
| DTP | 0.865 | [0.656, 1.000] | > 0.004 mg L<sup>-1</sup> | 0.75 | 1.00 |
| SRP | 0.862 | [0.686, 1.000] | > 0.004 mg L<sup>-1</sup> | 0.62 | 1.00 |
| Chl-*a* | 0.859 | [0.708, 0.974] | > 1.69 µg L<sup>-1</sup> | 0.88 | 0.79 |
| NO<sub>3</sub><sup>-</sup>+NO<sub>2</sub><sup>-</sup> | 0.849 | [0.566, 1.000] | > 0.401 mg L<sup>-1</sup> | 0.88 | 0.92 |
| **Multivariate (all 7)** | **0.974** | **[0.901, 1.000]** | — | — | — |

All seven single-variable classifiers achieved AUC values above 0.84. TP gave the highest AUC of 0.943 (95% bootstrap CI: 0.803–1.000), closely followed by chloride (0.924, [0.821, 0.995]) and turbidity (0.875, [0.724, 0.983]). The remaining four variables (DTP, SRP, Chl-*a*, and NO□□+NO□□) had AUC values between 0.849 and 0.865. The multivariate model combining all seven variables had an AUC of 0.974 (0.901–1.000) (Table 2).

Optimal decision thresholds identified by Youden’s J statistic were 0.030 mg L^-1^ for TP, 10.8 mg L^-1^ for chloride, 20.4 NTU for turbidity, 0.004 mg L^-1^ for DTP, 0.004 mg L^-1^ for SRP, 1.69 μg L□¹ for Chl-*a*, and 0.401 mg L^-1^ for NO□□+NO□□. At their respective optimal thresholds, TP achieved 88% bootstrap mean sensitivity and 100% specificity, chloride achieved 100% bootstrap mean sensitivity and 79% specificity, and turbidity achieved 88% bootstrap mean sensitivity and 88% specificity.

The optimal thresholds for TP, chloride, and turbidity were close to the inflection points of the partial dependence curves in Fig. 7b-e, where the two analyses are based on different statistical frameworks applied to the same dataset.

## 4. Discussion

### 4.1 Episodic *blaKPC* pulses linked to local precipitation events and wet-weather wastewater inputs

This study followed clinically relevant ARGs across a full year in the Detroit River, a binational channel that feeds downstream drinking-water supplies. For surveillance, what matters is not only whether these genes occur but when and from where they enter the river. *blaKPC* arrived as short spring pulses, whereas *mecA* persisted year-round, and these two patterns point to different sources and different surveillance implications.

*blaKPC* in the Detroit River was not characterized by its average concentration, but by its highly uneven distribution over time. A few spring weeks carried most of the annual signal, while levels stayed near zero for the rest of the year. This is the signature of a pulse-driven system. In contrast, *mecA* showed a more continuous pattern throughout the year. Three lines of evidence indicate that the *blaKPC* pulses were driven by local precipitation and wet-weather inputs from the urbanized sewershed, not by basin-scale runoff or snowmelt.

The first line of evidence is temporal. The year’s two largest rainfall events, in early April and early May, were each followed within a week by spring maxima in chloride, phosphorus, and dissolved nitrogen, and the early-May event coincided with the annual *blaKPC* peak (Fig. 2a, 4, S1). The earlier *blaKPC* peak on 28 April fell within the same wet spring period, although weekly composite sampling could not resolve its exact rainfall trigger. River discharge stayed flat from March through November with no spring rise (Fig. S1b). Because the Detroit River is a regulated connecting channel with flow controlled by upstream Great Lakes outflow (Scavia et al., 2019), the flat discharge indicates that no basin-scale runoff or snowmelt pulse contributed to the channel. Seasonal snowfall at Windsor in winter 2024-2025 was about 54 cm, well below the 1981-2021 normal of 129 cm, leaving little accumulated snow to melt in spring (ECCC, 2025).

The second line of evidence comes from wastewater normalization. The *blaKPC*/PMMoV ratio showed no significant seasonal variation, indicating that *blaKPC* and the wastewater tracer increased in parallel during the spring pulses. One reading is that the spring increase reflected a larger wastewater contribution rather than a higher *blaKPC* load per unit of sewage, though PMMoV is only an indirect proxy for wastewater input and the volume itself was not measured. Because the channel discharge was stable, any increase would more likely reflect local wet-weather inputs from the urbanized sewershed than basin-scale dilution. Detroit and Windsor both retain legacy combined sewer, and wet-weather overflows from them are a recognized contributor to nearshore water quality (Hartig et al., 2021; Li et al., 2003). Precipitation has been directly linked to increased ARG abundance in storm-drain outfalls and in combined sewer overflow discharges in comparable urban systems (Ahmed et al., 2018; Garner et al., 2017; Honda et al., 2020).

The third line of evidence is the suite of water-quality parameters that increased during the spring *blaKPC* pulse weeks. Chloride climbed to more than three times its summer and autumn level of approximately 9.4 mg L^-1^, too large a shift for routine sanitary sewage, which carries a smaller and steadier chloride signal (Corsi et al., 2015; Mackie et al., 2022). This pattern is more consistent with the flushing of accumulated road-salt residues from impervious urban surfaces into storm drains and combined sewers during spring rainfall, a mechanism documented in snow-affected urban watersheds even during mild winters (Corsi et al., 2015; Mackie et al., 2022). The accompanying turbidity peak fits mobilization of urban sediment by storm runoff (Brezonik and Stadelmann, 2002; Müller et al., 2020), and the simultaneous rise in TP and NO□□+NO□□matches the nutrient load of urban runoff, with a likely contribution from CSO discharge (Brezonik and Stadelmann, 2002; Long et al., 2014). The chemical signature of the *blaKPC* pulse weeks likely reflects a wet-weather urban runoff and sewer-overflow signal, rather than a baseline wastewater or basin-scale runoff signature.

Across these lines of evidence, the spring *blaKPC* pulses in the Detroit River are consistent with precipitation-associated wet-weather inputs from the local urbanized sewershed, including stormwater runoff and combined-sewer overflows. Annual averages would obscure these short windows, and ARG surveillance without targeted wet-weather sampling would miss most of the relevant exposure (Bengtsson-Palme et al., 2023; Pruden et al., 2021). *mecA* did not track these spring pulses, which is consistent with the two genes responding to different inputs over the year rather than to a single shared pathway.

### 4.2 Persistent *mecA* dynamics reflect distinct source mechanisms

Unlike the episodic *blaKPC* pulses, *mecA* persisted year-round, which points to a more continuous set of inputs rather than a single event-driven pathway. Several sources could sustain this background. *mecA*-carrying hosts may be widely distributed or well adapted to the river, and some inputs may not be scaled with the fecal tracer. The wastewater-normalized ratios offer one way to separate these possibilities.

Normalizing PMMoV separated the two genes more clearly than the raw concentrations did. The *blaKPC*/PMMoV ratio did not vary across seasons, whereas the *mecA*/PMMoV ratio did. The *mecA* ratio was lowest in spring, when the PMMoV signal was highest, and rose in summer and autumn as PMMoV declined. *mecA* and the fecal tracer therefore moved out of phase during much of the year. A purely wastewater-driven source would not produce this pattern, so part of the summer and autumn *mecA* signal likely came from inputs that PMMoV does not track.

One candidate is a pool of methicillin-resistant staphylococci that persists outside the human gut and wastewater pathway. *mecA* has been recovered from coagulase-negative staphylococci (CoNS) in surface waters, which suggests that environmental staphylococci can carry methicillin-resistance determinants that need not scale with sewage volume (Gómez et al., 2017; Larsson and Flach, 2022). PMMoV indexes enteric fecal input while staphylococci are largely skin-associated and environmental, an environmental CoNS pool could add *mecA* without a proportional increase in the fecal tracer. Waterborne staphylococci often originate from skin shedding rather than fecal pollution, and conventional fecal indicators do not reliably reflect their presence in water (Plano et al., 2011; Steadmon et al., 2024). The raw *mecA* signal stayed detectable year-round at both sites, while only the *mecA*/PMMoV ratio shifted seasonally.

Sediments may provide another pathway. River sediments accumulate ARG through continuous deposition and harbor bacterial densities that can exceed those in the overlying water column (Heß et al., 2018; Reichert et al., 2021). *mecA* has also been quantified in stream sediments at concentrations higher than those in the water column (Heß et al., 2018). Resuspension of these reservoirs by wind-driven turbulence, vessel traffic, or autumn mixing could re-introduce sediment-associated *mecA* gene or its microbial host into the water column without requiring a fresh wastewater input (Heß et al., 2018).The Detroit River nearshore zone is shallow, more exposed to wind and to ship wakes, and bordered by depositional fine sediments, all of which are conditions under which resuspension may become important in late summer and autumn. The autumn nearshore peak, which occurred without a corresponding PMMoV peak at that site, is consistent with this mechanism.

The nearshore-offshore contrast is also consistent with this interpretation. The nearshore site lies in the shallower, more urbanized portion of the channel and held higher autumn *mecA* than the offshore site over several weeks. Nearshore zones are more directly exposed to local non-wastewater inputs, including urban stormwater, shoreline runoff, wildlife and waterfowl fecal contamination, and resuspension from shoreline sediments. Stormwater has been shown to contribute ARG loads to urban streams, and sediments can act as important ARG reservoirs in wastewater treatment plant-impacted rivers (Garner et al., 2017; Reichert et al., 2021). The lateral pattern is consistent with the hydrodynamic structure of the Detroit River. The channel is generally well mixed vertically but not well mixed laterally, and transect studies have shown that nutrient concentrations close to shore can be routinely higher than those in the main central channel (Burniston et al., 2018; Scavia and Calappi, 2023). This lateral structure is large enough to shape where loads travel. In the lower Detroit River, the Amherstburg channel transports 37% of the discharge but a larger share of nutrient mass (Scavia and Calappi, 2023). A single sampling position therefore captures only one part of the river’s water-quality signal. Although lateral ARG transects are not available for the present dataset, the autumn divergence between nearshore and offshore *mecA* concentrations is consistent with this lateral structure, reflecting partly separate combinations of local sources and channel-scale mixing.

Overall, the persistent and weakly correlated *mecA* pattern is best read as a background signal fed by several overlapping sources rather than by one event. The year-round detection at both sites, the seasonal swing in the *mecA*/PMMoV ratio, and the candidate environmental and sediment reservoirs together point in this direction, in contrast to the more event-linked behavior of *blaKPC*. For surveillance, this means *mecA* is more useful as a persistent baseline indicator than as a marker of discrete contamination events. Although this study cannot resolve the dominant seasonal reservoir of *mecA*, its consistent detection across sampling contexts highlights its value as a baseline marker for longitudinal AMR surveillance.

### 4.3 The multivariate nature of AMR-water-quality relationships

The non-significant year-round Spearman correlation between *blaKPC* and individual water-quality variables can be reconciled with the strong pulse-week-stratified differences once the pulse-dominated character of the system is recognized. Because *blaKPC* stayed below detection or near zero for most weeks, the year-round correlation was dominated by a long low-signal baseline, and the few spring pulse weeks carried little weight in the stretches (Karkman et al., 2019). Hydrological events also drive concurrent changes in runoff, particulates, nutrients, chloride, and microbial inputs, so any single variable reflects only part of the underlying response (Almakki et al., 2019; Müller et al., 2020).

Pulse-week stratification, PCA, and random forest analysis recover this signal through three different routes. Stratification removes the quiescent baseline that diluted the annual statistics, and the contrast between pulse weeks and background weeks becomes detectable with large effect sizes (Garner et al., 2017; Honda et al., 2020). In the PCA, the water-quality variables that rose together during pulse weeks shared a common axis of variation, so most of the spring signal was carried by PC1 rather than spread across ten variables (Amos et al., 2015; Reichert et al., 2021). The random forest classifier captured the non-linear shape of the response. Predicted spring-week probability rose sharply once individual variables crossed narrow empirical ranges, and these inflection points fell close to the ROC-derived Youden’s thresholds obtained through a separate analysis (Jang et al., 2021; Jiang et al., 2024; Zahra et al., 2023).

### 4.4 Water-quality thresholds as early warning indicators for *blaKPC* risk

Routine qPCR and metagenomic surveillance of ARGs in surface water remain technically demanding and cost-intensive, which has limited its adoption by water utilities outside of research contexts (Cutrupi et al., 2024; Liguori et al., 2022; Pruden et al., 2021). Our results point to a more practical alternative. Conventional water-quality parameters already measured by existing regulatory and watershed monitoring programs performed well as proxy indicators for periods of elevated *blaKPC* risk in the Detroit River, with TP achieving an AUC of 0.943 and chloride an AUC of 0.924 (Table 2). These values are comparable to those reported in recent machine learning approaches to AMR surveillance in aquatic systems (Calero-Cáceres et al., 2026; Jang et al., 2021; Jiang et al., 2024; Sun et al., 2021; Zahra et al., 2023).

The optimal thresholds identified here (TP > 0.030 mg L^-1^, chloride > 10.8 mg L^-1^, turbidity > 20.4 NTU) fall within concentration ranges reported for spring runoff in Laurentian Great Lakes tributaries (Dove and Chapra, 2015; Hrycik et al., 2024; Mackie et al., 2022; Seilheimer et al., 2013). The overlap is consistent with the interpretation that wet-weather runoff drives the spring *blaKPC* pulses in this system. For management purposes, the existing binational monitoring networks along the Detroit River corridor are already set up to identify the conditions under which *blaKPC* risk peaks, without needing dedicated AMR sampling infrastructure.

The multivariate logistic model combining all seven parameters achieved the highest AUC of 0.974, exceeding the best single-variable model by 0.031. The bootstrap confidence intervals overlapped substantially (multivariate: 0.901–1.000; TP: 0.803–1.000), indicating that although the multivariate model performed better, the magnitude of this improvement should be interpreted cautiously. For operational deployment, a tiered monitoring framework in which exceedance of a single variable threshold triggers confirmatory *blaKPC* sampling may therefore capture most of the predictive signal while avoiding the computation and data integration costs of a full multivariate model (Huijbers et al., 2019; Pruden et al., 2021).

### 4.5 Transboundary surveillance and One Health implications

The Detroit River is a shared waterway between Canada and the United States, and its hydrological, chemical, and microbiological signals are inherently shared across jurisdictions (Beach et al., 2025; Corchis-Scott et al., 2024). A *blaKPC* pulse arriving at the offshore site is not constrained by political boundaries. The same water continues into Lake Erie, which supplies drinking water to more than 12 million people across Ontario, Michigan, Ohio, Pennsylvania, and New York (Reutter, 2019). The patterns reported here therefore have implications beyond a single river segment, and are consistent with the One Health framing of AMR (Larsson and Flach, 2022; McEwen and Collignon, 2018).

Two of the markers we tracked are clinically relevant. Carbapenem-resistant Enterobacterales carrying *blaKPC* are in the critical-priority tier of the WHO Bacterial Priority Pathogens List, and methicillin-resistant *Staphylococcus aureus* is in the high-priority tier (Sati et al., 2025). They were prioritized because resistance in these organisms strips away last-line treatment options, and carbapenem-resistant Enterobacterales infections in particular carry high mortality with few remaining alternatives (Munoz-Price et al., 2013). Their detection in spring runoff in a transboundary urban connecting channel does not in itself translate into clinical exposure risk, because gene copy concentrations are not a direct measure of viable, transmissible bacteria, and the pathway from environmental ARG to human carriage involves several poorly quantified steps (Bengtsson-Palme et al., 2023; Larsson and Flach, 2022). The data more clearly indicate that wastewater derived determinants of clinical relevance enter a shared aquatic environment through identifiable hydrological events, and that they do so for several weeks each year at concentrations well above background. This is consistent with a growing set of environmental compartments where seasonal AMR loading is now documented at watershed scale (Hendriksen et al., 2019; Pruden et al., 2021).

These findings also point to a practical route for transboundary surveillance. Our analysis used routinely collected water-quality variables from ECCC’s Great Lakes Connecting Channels Monitoring and Surveillance Program, which supports commitments under the Great Lakes Water Quality Agreement (Krantzberg, 2012; Scavia et al., 2019). Phosphorus, chloride, turbidity, and dissolved nitrogen were among the strongest predictors of spring *blaKPC* risk. Existing water-quality data streams could therefore be extended with an AMR risk-flag layer, so that spring-runoff conditions linked to elevated *blaKPC* risk trigger confirmatory ARG sampling across the corridor. This would integrate AMR surveillance into established water-quality monitoring systems rather than creating a parallel standalone program (Berendonk et al., 2015; Hendriksen et al., 2019).

Operationalizing this approach may require harmonization across agencies. Canadian and US programs may differ in sampling frequency, reporting periods, laboratory protocols, and unit conventions, and qPCR assays for ARGs are not yet routine in surface-water monitoring. Threshold values such as those identified here would be comparable across jurisdictions only if ARG assay panels, water-quality measurements, recovery controls, and data reporting were standardized under a shared framework (Bengtsson-Palme et al., 2023; Liguori et al., 2022).

Although this analysis is limited to a single watershed and year, the broader observation is testable in other cold-climate transboundary systems, including Great Lakes connecting channels, the St. Lawrence corridor, and European border rivers. Cross-watershed comparisons would help determine whether pulse-week-stratified water-quality thresholds can support a transferable approach to environmental AMR surveillance (Berendonk et al., 2015; Pruden et al., 2021).

### 4.6 Limitations and future directions

These findings show that clinically relevant ARGs followed different source pathways and that ARG-water-quality relationships were pulse-driven. However, the analysis was based on 10 months of weekly sampling in a single watershed, therefore the threshold values reported here should be treated as exploratory and specific to the Detroit River. The eight spring weeks used in the classification and ROC analyses are a small sample, which widened the bootstrap confidence intervals for several thresholds. Multi-year sampling would be needed to test whether these threshold ranges persist under different precipitation and runoff timing, wastewater infrastructure, and seasonal conditions (d’Orgeville et al., 2014; Hrycik et al., 2024; Reid et al., 2019). Water-quality measurements were available only at the offshore site. Paired lateral transects of AMR and water quality would help resolve whether the nearshore-offshore *mecA* divergence reflects distinct source pathways or channel-scale mixing (Scavia and Calappi, 2023). PMMoV provided a useful wastewater-associated normalization marker, but its behavior in flowing surface water can be affected by dilution, particle association, and environmental persistence (Kitajima et al., 2018; Symonds et al., 2019). Future work combining longer time series, and short- or long-read metagenomics could clarify the sources, hosts, and broader resistome associated with runoff events. Comparable analysis in other cold-climate transboundary watersheds would further show whether the pulse-weekt-stratified framework can support a more transferable approach to environmental AMR surveillance.

## 5. Conclusions

Year-round weekly monitoring of the Detroit River in 2025 showed clearly different temporal patterns for two clinically important resistance genes. *blaKPC* was pulse driven: one week accounted for 37.2% of the annual offshore signal, and the top three weeks for 72.8%. *mecA* was present in almost every sample with no dominant pulse. The *blaKPC*/PMMoV ratio stayed constant across seasons, consistent with larger wastewater volumes during spring wet-weather inputs rather than a new source. The *mecA*/PMMoV ratio was lowest in spring, pointing to non-wastewater inputs from environmental staphylococci or sediment reservoirs.

Spearman correlation between *blaKPC* and individual water quality variables were all non-significant. Pulse-week-stratified and multivariate analysis recovered the signal that linear statistics missed. Seven parameters were significantly elevated during spring *blaKPC* pulse weeks, PCA separated spring weeks from background, and a random forest classifier reached a leave-one-out AUC of 0.917. AMR water quality relationships in this system are non-linear and pulse dependent.

Conventional water quality variables performed well as proxies for spring *blaKPC* risk, single-variable AUCs exceeded 0.85 and the multivariate model reached 0.97. Optimal thresholds coincided with the inflection points of the partial dependence curves, so two independent frameworks converged on the same values. Existing binational monitoring along the corridor already records these variables, so an AMR risk flag layer could be added without new sampling infrastructure.

These thresholds are specific to one year and one watershed. Multi-year monitoring, additional indicator genes, and shotgun metagenomics will be needed to test transferability. Combining pulse-week-stratified analysis with routine water quality data offers a practical route for integrating AMR surveillance into established programs in cold-climate transboundary watersheds.

## Acknowledgement

We thank the Freshwater Quality Monitoring Division of Environment and Climate Change Canada (ECCC) for its support of the monitoring station (Environment and Climate Change Canada, 2025). We acknowledge research funding support through the WE SPARK Igniting Discovery Grants program held by OL, jointly funded by the Black Scholars Institute, the Great Lakes Institute for Environmental Research, the Faculty of Science (University of Windsor), and the WE-SPARK Health Institute, as well as funding provided by the Canada Research Chairs Program held by OL. Additional support was provided by INSPIRE (Integrated Network for the Surveillance of Pathogens: Increasing REsilience and capacity in Canada’s pandemic response), a program funded through the Canada Biomedical Research Fund Stage 2 (grant no. CBRF2-70 2023-00008), the Biosciences Research Infrastructure Fund Stage 2, and the Ontario Research Fund.

## Supplementary Material

**Fig. S1** Temporal correspondence between precipitation, Detroit River discharge, and offshore *blaKPC* pulses in 2025. (a) Daily and 7-day cumulative precipitation at Windsor Airport; major rainfall on 2 April and early May preceded the spring *blaKPC* pulses. (b) Detroit River discharge remained relatively stable from March to November, with no spring peak. (c) Offshore *blaKPC* maxima on 28 April and 5 May coincided with elevated 7-day precipitation, but not increased river discharge. Pink bands indicate spring weeks with *blaKPC* > 186 copies L□¹.

Table S1. Primers and probes used for qPCR and RT-qPCR assays in this study

Table S2. Reaction setup and thermal cycling conditions for qPCR and RT-qPCR assays used in this study

