## Supplementary figures and images for "Pulse-driven and persistent antimicrobial resistance markers in a transboundary Great Lakes connecting channel: pulse-week-stratified water-quality thresholds for One Health surveillance"

### Figure S1

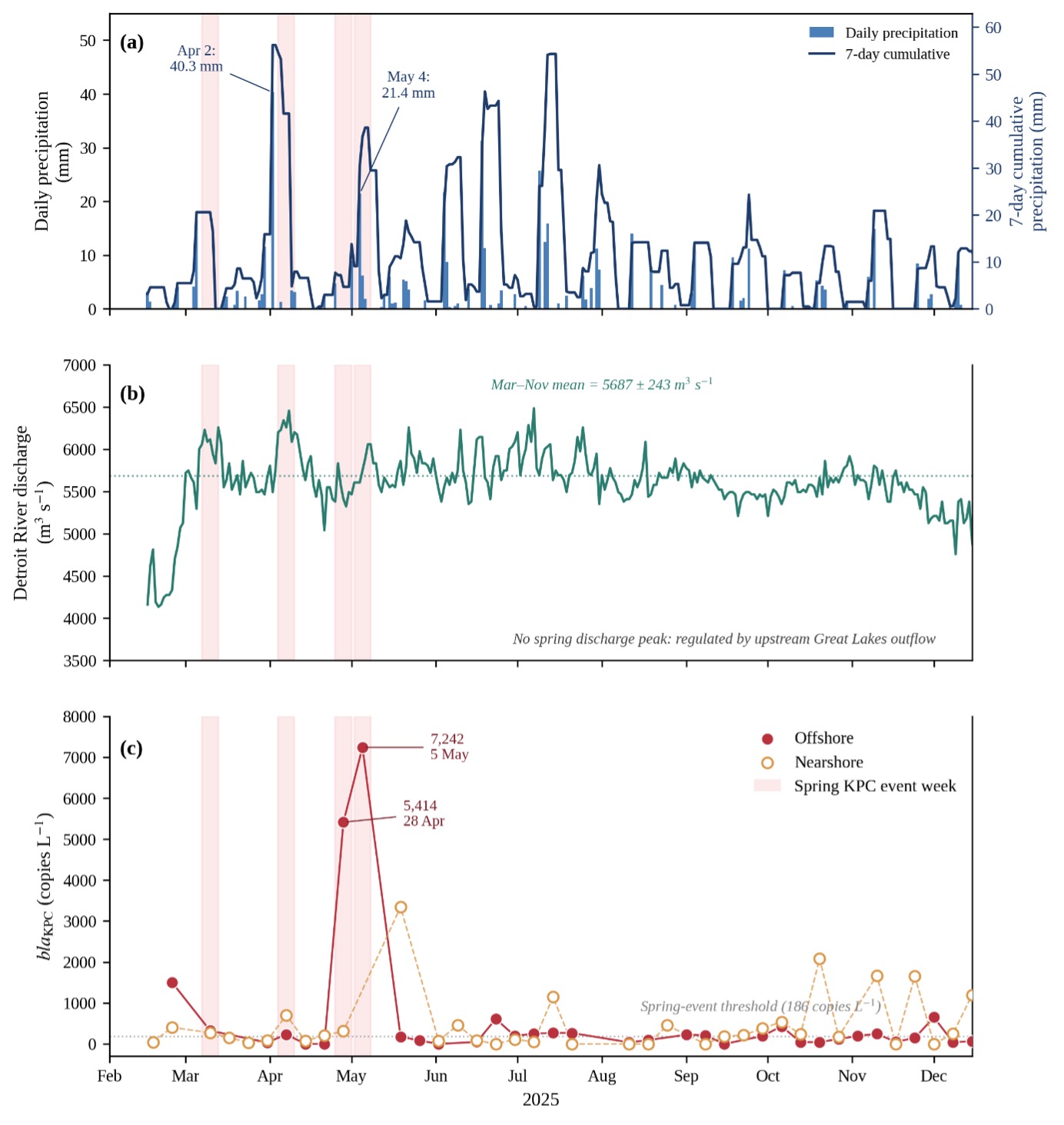
