## Supplementary material for "Pulse-driven and persistent antimicrobial resistance markers in a transboundary Great Lakes connecting channel: pulse-week-stratified water-quality thresholds for One Health surveillance": Table S1

**Table S1. Primers and probes used for qPCR and RT-qPCR assays in this study.**

| **Target** | **Forward primer (5′–3′)** | **Reverse primer (5′–3′)** | **Probe (5′–3′)** | **Final conc. (nM)** | **Amplicon length** | **Reference** |
| --- | --- | --- | --- | --- | --- | --- |
| *blaKPC* | GGCCGCCGTGCAATAC | GCCGCCCAACTCCTTCA | FAM-TGATAACGCCGCCGCCAATTTGT | Primers 800  Probe 400 | 61 bp | Hindiyeh et al., 2008; Lee et al., 2015 |
| *blaNDM* | GACCGCCCAGATCCTCAA | CGCGACCGGCAGGTT | HEX-TGGATCAAGCAGGAGAT | Primers 800  Probe 400 | 52 bp | Lee et al., 2015; Bordin et al., 2019 |
| *blaVIM-2* | AATGGTCTCATTGTCCGTGATG | TACAGCGTGGGGTGCGA | Cy5-TGATGAGTTGCTTTTGATTG | Primers 800  Probe 400 | 61 bp | Brown-Jaque et al., 2018 |
| *mecA* | CAATGCCAAAATCTCAGGTAAAGTG | AACCATCGTTACGGATTGCTTC | HEX-ATGAGCTATATGAGAACGG | Primers 900  Probe 250 | 107 bp | Galia et al., 2019 |
| *mcr-1* | CTGGGCGCGGATGAGTAT | AGCGTATCHAGCACATTTTCTTG | FAM-ATGTCGATACCGCCAAA | Primers 200  Probe 100 | 68 bp | Gong et al., 2024 |
| PMMoV | GAGTGGTTTGACCTTAACGTTTGA | TTGTCGGTTGCAATGCAAGT | Cy5-CCTACCGAAGCAAATG | Primers 200  Probe 200 | 68 bp | Zhang et al., 2006; Rosario et al., 2009; Haramoto et al., 2013 |

**Note:** All primers and probes were obtained from Integrated DNA Technologies (IDT, Coralville, IA, USA).
