## Supplementary material for "Pulse-driven and persistent antimicrobial resistance markers in a transboundary Great Lakes connecting channel: pulse-week-stratified water-quality thresholds for One Health surveillance": Table S2

**Table S2. Reaction setup and thermal cycling conditions for qPCR and RT-qPCR assays used in this study.**

| **Assay** | **Reaction chemistry** | **Reaction setup (20 μL final volume)** | **Thermal cycling conditions** |
| --- | --- | --- | --- |
| *blaKPC/blaNDM/blaVIM-2* triplex qPCR | Luna Universal Probe qPCR Master Mix | 10 μL 2× master mix, primers and probes at final concentrations listed in Table S1, 5 μL template, and nuclease-free water to 20 μL | 95 °C for 20 s; 45 cycles of 95 °C for 5 s and 55 °C for 30 s |
| *mcr-1* qPCR | Luna Universal Probe qPCR Master Mix | 10 μL 2× master mix, primers and probe at final concentrations listed in Table S1, 5 μL template, and nuclease-free water to 20 μL | 25 °C for 1 min; 95 °C for 5 min; 45 cycles of 95 °C for 15 s and 55 °C for 30 s |
| *mecA* qPCR | Luna Universal Probe qPCR Master Mix | 10 μL 2× master mix, primers and probe at final concentrations listed in Table S1, 5 μL template, and nuclease-free water to 20 μL | 25 °C for 1 min; 95 °C for 5 min; 45 cycles of 95 °C for 15 s and 60 °C for 30 s |
| PMMoV RT-qPCR | Luna Universal Probe One-Step Reaction Mix with Luna WarmStart RT Enzyme Mix | 10 μL 2× reaction mix, 1 μL RT enzyme mix, primers and probe at final concentrations listed in Table S1, 5 μL template, and nuclease-free water to 20 μL | 55 °C for 10 min; 95 °C for 1 min; 40 cycles of 95 °C for 10 s and 55 °C for 30 s |
